# The opportunity cost of time modulates cognitive effort

**DOI:** 10.1101/201863

**Authors:** A. Ross Otto, Nathaniel D. Daw

**Affiliations:** Department of Psychology, McGill University; Princeton Neuroscience Institute and Department of Psychology, Princeton University

## Abstract

A spate of recent work demonstrates that humans seek to avoid the expenditure of cognitive effort, much like physical effort or economic resources. Less is clear, however, about the circumstances dictating how and when people decide to expend cognitive effort. Here we adopt a popular theory of opportunity costs and response vigor and to elucidate this question. This account, grounded in Reinforcement Learning, formalizes a trade-off between two costs: the harder work assumed necessary to emit faster actions and the opportunity cost inherent in acting more slowly (i.e., the delay that results to the next reward and subsequent rewards). Recent work reveals that the opportunity cost of time—operationalized as the average reward rate per unit time, theorized to be signaled by tonic dopamine levels, modulates the speed with which a person responds in a simple discrimination tasks. We extend this framework to cognitive effort in a diverse range of cognitive tasks, for which 1) the amount of cognitive effort demanded from the task varies from trial to trial and 2) the putative expenditure of cognitive effort holds measureable consequences in terms of accuracy and response time. In the domains of cognitive control, perceptual decision-making, and task-switching, we found that subjects tuned their level of effort exertion in accordance with the experienced average reward rate: when the opportunity cost of time was high, subjects made more errors and responded more quickly, which we interpret as a withdrawal of cognitive effort. That is, expenditure of cognitive effort appeared to be modulated by the opportunity cost of time. Further, and consistent with our account, the strength of this modulation was predicted by individual differences in efficacy of cognitive control. Taken together, our results elucidate the circumstances dictating how and when people expend cognitive effort.

## Introduction

“There is no expedient to which a man will not resort to avoid the real labor of thinking.”

-- Sir Joshua Reynolds

Many human behaviors necessitate a trade-off between effort and reward: how a person performs in a given task is due, in part, to his or her decision to expend cognitive effort in the service of obtaining rewards. A critical constraint underlying this trade-off is the inherently limited information-processing of the “central executive”, (Navon & Gopher, 1979; Norman & Shallice, 1986), which necessitates that cognitive processing resources be allocated in accordance with our behavioral goals. Accordingly, the question of when and why people decide to expend—or withhold—cognitive effort has been the subject of vigorous examination in recent literature (Boksem & Tops, 2008; Boureau, Sokol-Hessner, & Daw, 2015; Inzlicht, Schmeichel, & Macrae, 2014; Kool, McGuire, Rosen, & Botvinick, 2010; Kurzban, Duckworth, Kable, & Myers, 2013; Westbrook & Braver, 2016).

Much of this work takes the perspective that the level of processing adopted by an individual at a given moment—in, say, a response conflict task—is chosen strategically on the basis of a cost-benefit analysis (Botvinick & Braver, 2015; Boureau et al., 2015; Gratton, Coles, & Donchin, 1992; Sandra & Otto, 2018; Shenhav et al., 2017). However, although people consistently and systematically appear to avoid cognitive demands (Dreisbach & Fischer, 2012; Kool et al., 2010; Schouppe, Demanet, Boehler, Ridderinkhof, & Notebaert, 2014; Westbrook, Kester, & Braver, 2013), a key difficulty in achieving a decision-theoretic understanding of mental effort allocation is quantifying, or even defining, the benefits and especially the costs of cognitively effortful strategies. Highlighting this question, a number of lines of research find that even when cognitive demands are held constant over time, an individual’s exertion of flexible control over behavior fluctuates considerably over time—and these shifts thought to arise—in part—from changes in the perceived costs and benefits of engaging the central executive (Botvinick & Braver, 2015; Braver, Reynolds, & Donaldson, 2003; Kahneman, 1973). Although the nature of cognitive costs (for instance, whether or not cognitive effort depletes some objective resource, like energy) is unclear, we here focus on one subtype of cost – opportunity cost – whose existence and properties can be inferred from first principles. In this respect, a key, but untested, proposal is that internal cost signals represent, in whole or part, *opportunity costs*, owing to the limited nature of cognitive resources—that is, occupying cognitive resources in the service of a particular goal, for a particular time, foregoes the benefits that could be obtained by using them for some other goal (Botvinick & Braver, 2015; Kurzban et al., 2013; Shenhav et al., 2017). Up to now, there is little empirical evidence supporting the idea that an individual’s moment-to-moment allocation of cognitive resources are directed by opportunity costs. Here we provide a novel test of this idea, revealing how the outlay of cognitive effort across three diverse task paradigms is modulated by the opportunity cost of time.

To do this, we leverage an influential model of opportunity costs rooted in reinforcement learning (Niv, Daw, Joel, & Dayan, 2007), which has been previously applied to physical effort, to investigate the effects of opportunity costs on the allocation of cognitive effort. In its original formulation, this model formalizes a trade-off between two costs: the harder physical work assumed necessary to emit faster actions (“vigor”) and the opportunity cost inherent in acting more slowly. Thus, the opportunity cost of time—which in many settings equals the long-run average reward rate of the environment—should dictate response speed: when delayed action is more expensive, actions should be made more quickly because more rewards would be foregone by slow responses (Niv et al., 2007). Supporting this idea empirically, Guitart-Masip et al. (2011) and Beierholm et al. (2013) demonstrate how people adjust their response speeds in simple detection tasks depending on the prevailing reward rate, in accordance with the opportunity cost of time (Beierholm et al., 2013; Guitart-Masip, Beierholm, Dolan, Duzel, & Dayan, 2011). However, the reaction-time tasks employed by these studies were, by design, too simple to detect changes in response accuracy, or its relationship with response speed.

In a series of experiments, we test the idea that reliance upon cognitively demanding strategies should analogously shift with changing opportunity costs: when time is perceived to be expensive, we should shift our effort engagement so as to use quicker and less accurate, presumably cognitively inexpensive strategies to make decisions or solve tasks (Westbrook & Braver, 2016). Here, we draw upon Niv et al.’s formalization of response vigor to examine whether increasing the opportunity cost of time prompts a withdrawal of cognitive effort. Just as in the case of physical vigor, if the long-run average reward rate is higher in a cognitive task, then more reward can be obtained by moving on to the next trial sooner, which should shift the reward-rate-optimizing solution to accept a higher error rate (thus lower reward in a trial) by finishing more quickly. Such an effect, due to the opportunity cost of time, is predicted whether or not we assume that more or less cognitive engagement also carries additional, objective (e.g., energetic) costs. Following this computational framework and the empirical work it stemmed (Beierholm et al., 2013, 2013), we manipulated the average reward rate of the environment—and thereby the cost, in terms of foregone potential rewards, for acting slowly—while participants performed three distinct, but well-characterized cognitive tasks. For example, in simple response conflict paradigms, successfully overriding inappropriate, stimulus-driven responses requires engagement of effortful, ‘top-down’ control (Westbrook & Braver, 2015). Critically, in the tasks considered here, withholding cognitive effort (e.g., responding faster) carries a performance consequence in terms of response accuracy (and consequently, obtained rewards).

Accordingly, we reasoned that opportunity cost-evoked withdrawal of effort investment could manifest in simultaneous changes to speed and accuracy because disengagement of cognitive control is thought to decrease the quality of information accumulation (Cohen, Dunbar, & McClelland, 1990; Egner & Hirsch, 2005). At the same time, as the opportunity cost of time is demonstrated to speed responding overall (Beierholm et al., 2013; Guitart-Masip et al., 2011). Given that, in many cognitive task domains, fast responses incur larger error rates—a phenomenon known as the speed-accuracy tradeoff—a further important question is to what extent the pattern of changes we observe arises from some sort of strategic resource optimization against a fixed speed-accuracy tradeoff profile inherent to the task. An alternative possibility— also suggested by Niv et al. (2007) for the case of physical vigor—is that the general logic of opportunity costs is hardwired as a sort of reflexive or Pavlovian strategy, which automatically evokes faster responding in high average reward circumstances; even when this would ultimately be counterproductive (Boureau & Dayan, 2011). Across our three experiments we find some hints supporting for the latter perspective. First, we find that higher opportunity costs evoke faster, less accurate performance over and above what can be explained from the fixed speed-accuracy tradeoff as deduced at a fixed level of average reward. Second, we see analogous reward-related changes even in a task for which slower responding does not (holding average reward fixed) confer any accuracy benefit. We tentatively interpret these effects as reflecting more or less investment of cognitive effort, per se, so as to change (rather than merely optimize along) a fixed, task-imposed speed-accuracy tradeoff.

## Experiment 1: Perceptual Decision Task

We first sought to examine how the opportunity cost of time modulates the accumulation of information in a perceptual decision task, which itself costly in terms of time and presumably cognitive effort (Drugowitsch, Moreno-Bote, Churchland, Shadlen, & Pouget, 2012). Indeed, explicit time pressure alters the effort-accuracy tradeoff in multi-attribute and perceptual decision-making (Forstmann, Dutilh, et al., 2008; Payne, 1982) such that perceived time scarcity engenders decision strategies that utilize less information in order to make faster decisions.

### Methods

#### Participants

We recruited 37 participants on Amazon Mechanical Turk (AMT), an online crowdsourcing tool, to perform the two-alternative forced choice perceptual decision-making task. AMT allows experimenters to post small jobs to be performed anonymously by “workers” for a small amount of compensation (Crump, McDonnell, & Gureckis, 2013). Participants were all US residents, paid a fixed amount ($2 USD) plus a bonus contingent on their decision task performance, ranging from $1-3 USD. We excluded the data of 5 subjects who missed more than 20 response deadlines in either the calibration phase or the main task, yielding 32 subjects remaining in our subsequent analyses. The protocol for all the experiments was approved by the committee on human subjects at NYU, where the authors were affiliated when the experiments were conducted.

#### Calibration Phase

Before performing the main perceptual task, each subject underwent a calibration session to determine the stimulus set used in the main 2AFC task. Each trial, two squares were presented on screen, each of which were filled with 10×10 array of asterisks (*) in a 40-point font. The “reference” square always contained 50 dots, while the “variable” square contained of 20, 39, 43, 47, 53, 57, 61, or 80 dots (Figure 1A). Dots were placed randomly in the array for each square on each trial. The order of stimuli was randomly determined and evenly distributed except the extreme stimuli (20 and 80 points), which occurred on 3.2% of trials. Each trial, the two boxes were displayed simultaneously, and subjects had 600ms to choose which of the two squares contained more dots using the ‘E’ and ‘I’ buttons on the keyboard. To encourage quick responding 10% of these trials had deadlines of 500ms (Guitart-Masip et al., 2011). The stimuli remained on screen for 500ms following a response, and then feedback (“CORRECT” or “WRONG”) was displayed on screen for 1000ms. If no response was detected, the message “TOO SLOW” was displayed for 1000ms. Following feedback, a fixation cross was displayed with an inter-trial interval (ITI) ranging from 750 to 1250ms (uniformly distributed). Subjects completed 125 of these calibration trials.

**Figure 1.**
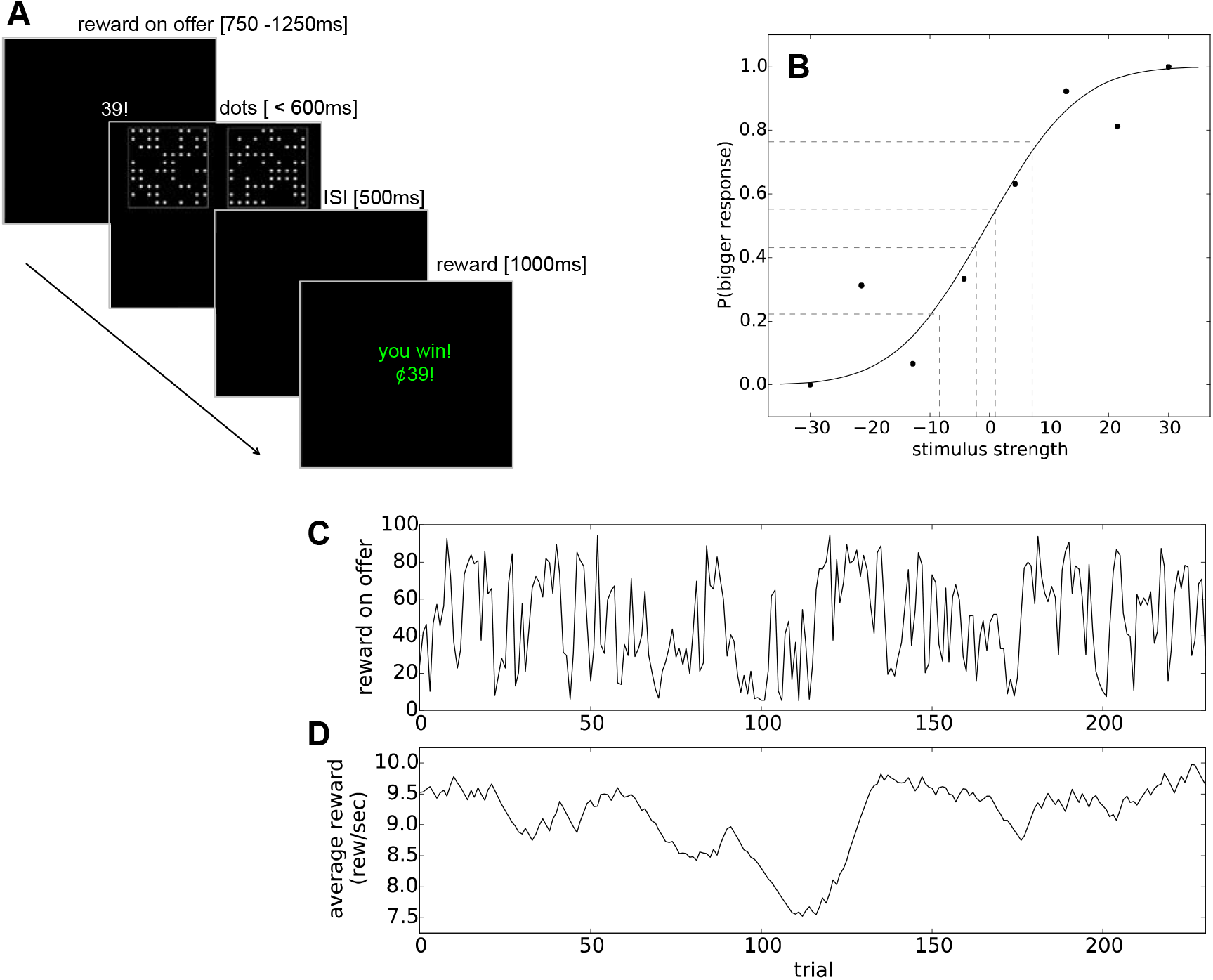
(A) Task flow in the perceptual decision-making experiment. Before the stimulus is displayed, subjects are shown the potential reward for making a correct response, then a decision is made about which square contains more dots, and following an inter-stimulus interval, the reward obtained is displayed. (B) An example subject’s data from the calibration phase of the perceptual decision-making experiment. Closed circles reflect the proportion of that the subjected indicated that the variable stimulus contained more dots as a function of stimulus strength. Lines indicate the fit of a cumulative Gaussian to these data. Dotted vertical lines indicate the stimulus strength values selected for use in the main experiment for this subject. (C) Top: we induced fluctuation in trial-to-trial available rewards (top), which, in conjunction with the subject’s history of responses, yielded an empirical average reward. Bottom: an example subject’s experienced average reward rate, in units of reward per second.

Stimulus sets used in the main phase were individually determined for each subject from online fits to the calibration data. Specifically, we identified stimulus strengths that led to “bigger” response probabilities of 0.23, 0.44, 0.56, 0.77, and 0.95 by fitting a cumulative normal psychometric function to each participants’ response data (Fleming, Maloney, & Daw, 2013; see Figure 1B). We recovered two parameters for each subjects: the mean of the cumulative normal distribution, which can be interpreted as response bias, and the standard deviation of the distribution, interpreted as the slope of the psychometric function. The average best-fitting values of bias and slope parameters were 0.56 (*SD*=1.48) and 11.69 (*SD*=7.05), respectively.

#### Main Phase

For each subject, dot stimuli strengths were randomly and uniformly sampled from the stimulus strengths selected in the calibration phase. Rewards available were determined randomly using a Gaussian random walk with standard deviation 30 and reflecting boundaries at 5 and 95 cents (Figure 1C). At the outset of each trial, subjects were presented visually with a number representing the reward on offer that trial, ranging from 1–100 cents (Figure 1A), which lasted from 750-1000ms, after which the same trial timing was used as in the calibration phase, with the exception that on correct trials, the feedback displayed was the reward obtained (e.g. “you win 9¢”) for 1000ms. Subjects completed as many dots trials as they could in the time limit of 20 min. Trials completed ranged from 152 to 312 trials (M=220). After completing the main phase, subjects were then paid a bonus proportional to their earnings in the task.

#### Regression Analyses

In the following analyses, focusing on the main phase of the experiment, we excluded the first 10 trials in order to allow subjects to acclimate to the procedure and excluded outlier trials with RTs greater than 3 standard deviations from each subject’s mean RT (< 2% of trials for all subjects). Following previous work we calculated average reward, 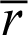, in units of reward/sec, using the following update rule (Constantino & Daw, 2015):

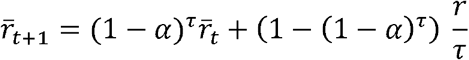

where r is the obtained reward on trial τ, is the time elapsed since the last update (which depends, critically, on each trial’s RT and ITI), and *α* is a learning rate parameter. Following previous work (Beierholm et al., 2013; Guitart-Masip et al., 2011) we fit a single learning rate to the RTs of the entire sample of subjects using a nonlinear optimization routine. However, rather than combine all subjects’ RTs and run a single regression, we accomplished this by running a separate regression for each subject finding the learning rate that minimizes the total error across the group. Specifically, the subject-level RT regression included the following terms:

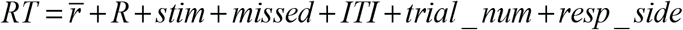

where RTs were log-transformed and z-scored, *R* is the reward available that trial, *stim* is the dots stimulus strength (expressed as the absolute value of the difference between the reference stimulus and the variable stimulus), *missed* is a binary variable representing whether the subject missed the previous trial’s response deadline, *trial_num* is a linear term representing trial number (to capture speeding over time), and *resp_side* represents whether a left or right-hand response was made (to capture simple response bias).

We found a best-fitting *α* of 0.0031. Note that this learning rate is smaller than that estimated in previous work (Beierholm et al., 2013) because the learning rule used here calculates average reward in unit of rewards per second rather than rewards per trial. We found that this formulation of average reward per unit time— which accords closely with the theoretical work on average reward and vigor (Niv et al., 2007)—explains RT variance considerably better than the average reward per trial.

To assess the influence of average reward upon RTs at the group level, we conducted mixed-effects regressions using the lme4 package (Pinheiro & Bates, 2000) in the R programming language, using the same predictors as the individual-level regression outlined above. All terms estimated at the fixed-effects level and as random effects at the subject level. Continuously-valued variables were inputted as within-subject z-scores. To examine accuracy we estimated a logistic regression with the same predictors but with the subject’s response (correct/error) as the outcome variable each. Significance values were computed using the car package in R (Fox & Weisberg, 2011).

Because the average reward predictor variable is determined by the free parameter *α*, which was itself estimated using a regression (resulting in the loss of a degree of freedom), it was critical to demonstrate that the regression results were not biased by the free parameter. To do this, we employed a cross-validation procedure whereby we fit the learning rate (using the procedure described above) to one half of the subjects, finding a best-fitting learning rate comparable to the full sample (*α* = 0.0026), and performed the regressions (as described above) on the remaining half the subjects. We recovered significant effects of average reward on both RT (*β*=-0.015, SE=0.0069, p=0.026) and accuracy *β*=-0.221, SE=0.0069, *p*<0.001)—mirroring the main results—suggesting that the estimation of the learning rate does not bias our estimation of average reward effects.

#### Drift Diffusion Model

We used hierarchical Bayesian estimation of drift diffusion model parameters, which has the advantage that fits to individual subjects are constrained by the group-level distribution (Frank et al., 2015; Wiecki, Sofer, & Frank, 2013). Hierarchical drift diffusion models (HDDMs) are particularly useful for estimating decision parameters—via regressions within the hierarchical model—that are allowed to vary from one trial to the next as a function of psychological or neural variables that vary from trial-to-trial. In particular, our regression was specified such that on each trial t, the threshold a and the drift rate *v* are influenced by both average reward (as calculated above) and reward on offer *R*:

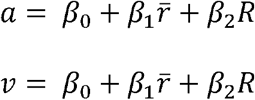

Estimation of the joint distribution of the parameters of the HDDM was performed using the hierarchical Bayesian implementation of the HDDM toolbox (version 0.6.1) via Markov Chain Monte Carlo techniques. Following previous work (Frank et al., 2015), each DDM parameter for each subject and condition was modeled to be distributed according to a normal (for real-valued parameters), or a Gamma (for positive-valued parameters), centered around the group mean with group variance. Prior distributions for each parameter were informed by several studies reporting best-fitting DDM parameters recovered on a range of decision-making tasks (Wiecki et al., 2013). Ten thousand samples were drawn from this model, discarding the first 2,000 samples for ‘burn-in.’

### Results

In the perceptual decision task, subjects made a series of judgments about which of two squares contained more dots (Figure 1A), following the task design of Fleming et al. (2014). Importantly, one square always contained 50 (out of 100 possible) dots, while the other square contained a variable amount of dots, which allowed us to examine subjects’ accuracy as a function of relative stimulus strength—here, the difference in number of dots between the two squares (Figure 1B). As subjects were free to make responses as quickly or as slowly as they wanted (up to a deadline), they could control the amount of evidence that they used to make a perceptual decision. Following a calibration phase, which ensured that test stimuli generated comparable accuracy levels across subjects (Fleming et al., 2013), subjects then made a series of perceptual decisions for the chance of receiving a monetary reward whose magnitude was shown at the beginning of each trial (Figure 1A).

To manipulate the perceived opportunity cost of time, we induced random fluctuations in these available rewards, which—in conjunction with an individual subject’s history of response times (RTs) and errors—yields a time-varying empirical average reward rate per second (Figure 1 C and D). As participants had a fixed amount of time in which to complete as many trials as possible, the prevailing average reward rate effectively imposes an “opportunity cost of time.” Thus, when the average reward rate is high (and thus, the opportunity cost of time is high), we expected participants to make faster responses, following previous findings (Beierholm et al., 2013; Guitart-Masip et al., 2011)—but at the expense of accuracy because less perceptual evidence can be accumulated during fast responses.

We then examined perceptual decision-making accuracy and response times (RTs) as a function of opportunity costs using a tertile split on experienced average reward rate, further grouping trials by difficulty of perceptual discrimination based on stimulus strength (yielding “easy” and “difficult” trials). We found that when the average reward rate was high, subjects made less accurate (Figure 2A) and faster (Figure 2B) responses compares to when the average reward rate was low. Following (Guitart-Masip et al., 2011) we estimated mixed-effects regressions (with a number of other predictor variables including reward currently ‘on offer’ and stimulus strength) to quantify the continuous effect of average reward rate upon accuracy and RTs (coefficient estimates reported in Tables 1 and 2). While reward ‘on offer’ exerted no significant effects on RT or accuracy, the average reward rate significantly sped RTs (*β*=-0.009, SE=0.004, *p*<.05) —corroborating previous findings (Beierholm et al., 2013; Guitart-Masip et al., 2011)—and here, also significantly reduced accuracy (*β*=-0.144, SE=0.033, *p*<.0001; Figures 2C and D).

**Figure 2.**
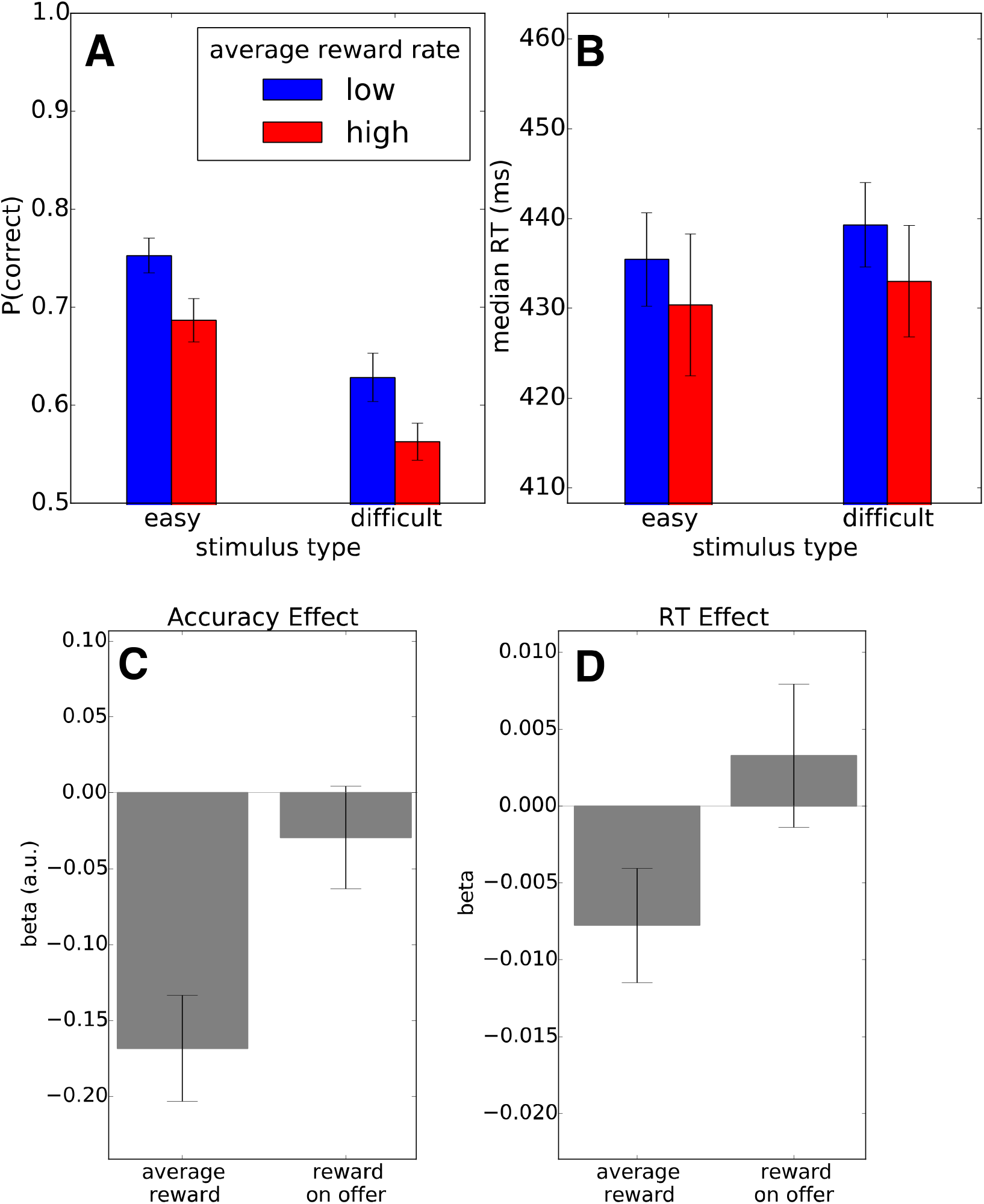
(A-B) Examining accuracy on easy trials (high stimulus strength) versus hard trials (low stimulus strength) as a function of the opportunity cost of time (lower versus upper tertile of average reward rate) revealed that responses were faster and less accurate when the opportunity cost of time was high. Error bars indicate standard error of the mean. (C-D) Mixed-effects regressions revealed that the average reward rate significantly reduced accuracy and sped RTs, but the amount of reward on offer on the present trial did not exert an effect on either RT or accuracy. Error bars indicate standard error of the regression coefficient estimate.

**Table 1:** Mixed-effects Regression coefficients indicating the influence of the average reward rate and a number of other trial-by-trial covariates upon RTs in Experiment 1 (Perceptual Decision-Making). Asterisks denote significance at the .05 level.

| *Coefficient* | *Estimate (SE)* | *p-value* |
| --- | --- | --- |
| (Intercept) | 6.070 (0.020) | <.0001* |
| iti | -0.013 (0.004) | <.0001* |
| **avg_reward** | **-0.009 (0.004)** | **0.010\*** |
| reward | 0.005 (0.004) | 0.350 |
| dots_difference | -0.004 (0.003) | 0.827 |
| prev_error | -0.018 (0.006) | <.0001* |
| trial_num | -0.018 (0.010) | 0.118 |
| resp_side | 0.006 (0.016) | 0.108 |

**Table 2:** Mixed-effects logistic regression coefficients indicating the influence of the average reward rate and a number of other trial-by-trial covariates upon accuracy in Experiment 1 (Perceptual Decision-Making). Asterisks denote significance at the .05 level.

| *Coefficient* | *Estimate (SE)* | *p-value* |
| --- | --- | --- |
| (Intercept) | 0.608 (0.087) | <.0001* |
| **avg_reward** | **-0.144 (0.033)** | **<.0001\*** |
| reward | -0.014 (0.028) | 0.601 |
| dots_difference | 0.751 (0.105) | <.0001* |
| prev_error | 0.021 (0.031) | 0.494 |
| trial_num | -0.001 (0.001) | 0.060 |
| resp_side | 0.037 (0.033) | 0.260 |

To elucidate in greater detail how opportunity costs influence task behavior, we jointly fit subjects’ choices and RTs with a drift diffusion model (DDM), a widely employed mathematical model of evidence accumulation, which has successfully explained choice behavior (both accuracy and RT) and neurophysiological measures in a variety of perceptual decision tasks (Gold & Shadlen, 2007; Ratcliff & Rouder, 1998). DDMs assume that evidence for one response over the other accumulates over time until the integrated evidence passes a threshold and a choice is made. The threshold parameter governs the speed-accuracy tradeoff between the benefits of accumulating more information with the cost of taking more time to reach a decision; we therefore hypothesized that the a high average reward would decrease the threshold of evidence accumulation. The DDM also allows us to explore whether evidence threshold changes are simultaneously accompanied by a change in quality of information accumulated (Rae, Heathcote, Donkin, Averell, & Brown, 2014).

Using hierarchical Bayesian model-fitting (Wiecki et al., 2013), we quantified, on a trial-by-trial basis, how the opportunity cost of time alters a subjects’ decision threshold. We found that the average reward rate, but not reward ‘on offer’ or stimulus strength, significantly reduced decision thresholds (Figure 3A)—that is, as average reward rate increased, responses were more likely to have a faster, more skewed RT distribution and have a higher probability of erroneous response, which is corroborated by the RTs and accuracies we found (Figures 2A and B). Indeed, the hypothesis that people should set evidence thresholds in accordance with the average reward rate has been difficult to demonstrate experimentally, by manipulating trial timing (i.e. delays; (Bogacz, Hu, Holmes, & Cohen, 2010). Here, people strategically modulate their evidence thresholds on a trial-to-trial basis in accordance with average reward rate, demonstrating the potency of this average reward manipulation.

**Figure 3.**
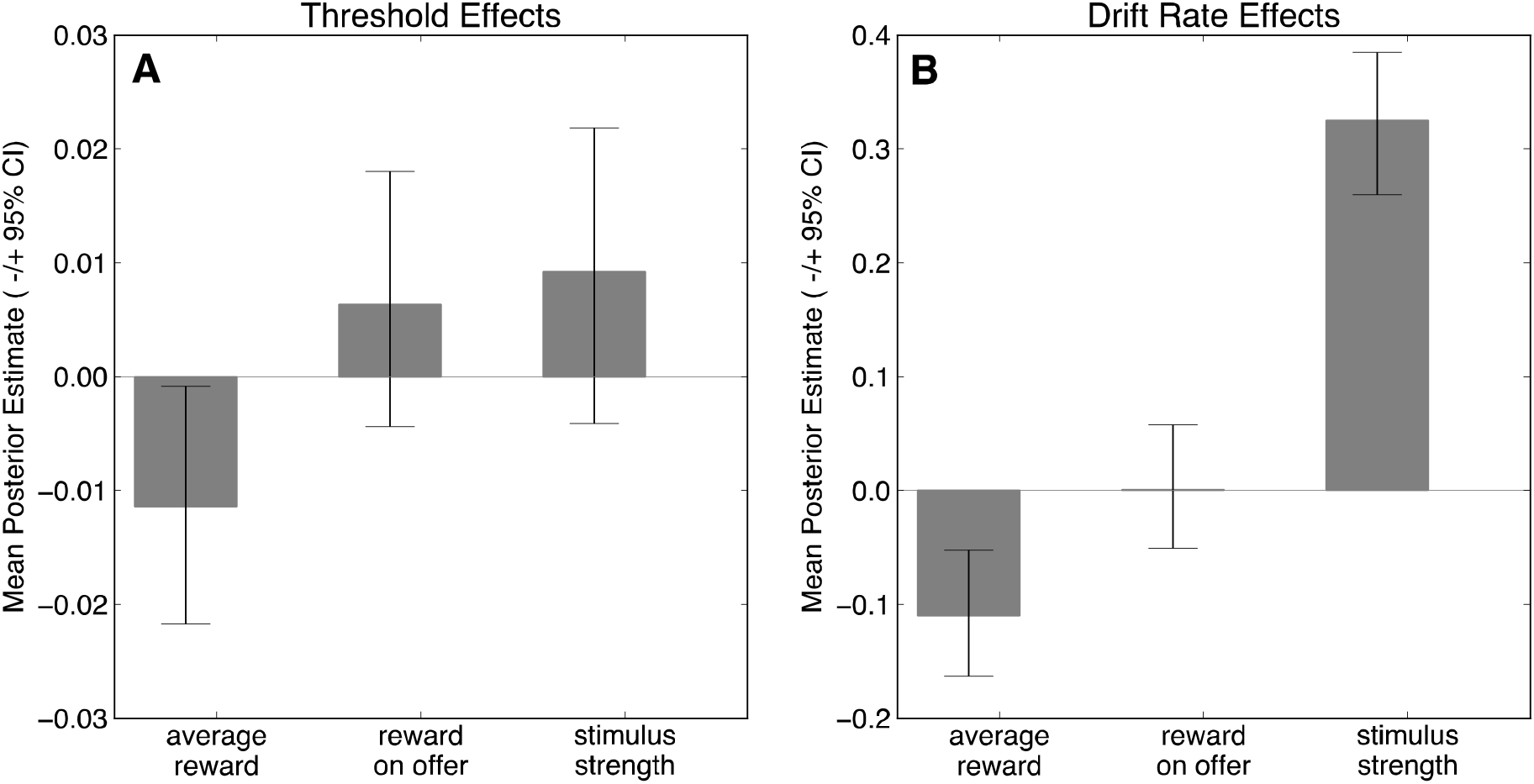
Posterior distributions of DDM parameters reveal that the average reward rate decreased decision thresholds (A), while drift rate decreased with the average reward rate and increased with strength of stimulus (B). Available reward did not affect either decision thresholds or drift rates. Error bars represent the 95% confidence interval of the posterior distribution.

A second parameter of the model controls the rate of evidence accumulation or “drift rate.” This parameter normally captures factors such as objective signal strength (Palmer, Huk, & Shadlen, 2005)—here, the difference in the number of dots between the two stimuli—and indeed increased with stimulus strength here (Figure 3B). Unexpectedly, we found that the drift rate was also negatively influenced by the average reward rate, suggesting that the opportunity cost of time did not solely induce a threshold adjustment (e.g., a change along a fixed speed-accuracy tradeoff), but could reflect a shift in the way evidence is processed (Dambacher & Hübner, 2015). This might represent a decreased investment of cognitive effort per se when opportunity costs are high, over and above the change in behavior attributable to decreased time investment (i.e., lower threshold), in line with the idea that evidence accumulation is itself cognitively effortful (Drugowitsch et al., 2012; Mathias et al., 2017).

In a subsequent experiment we more directly demonstrate that cognitive effort—beyond accumulation of perceptual evidence—can be modulated by these opportunity costs, using a classic cognitive control task (Forstmann, van den Wildenberg, & Ridderinkhof, 2008; Simon, 1990) for which participants need to inhibit pre-potent responses to respond accurately.

## Experiment 2: Simon Task

### Methods

#### Participants

We recruited 69 participants on AMT who paid a fixed amount ($2 USD) plus a bonus contingent on their decision task performance, ranging from $1-3 USD. We excluded the data of 19 subjects who missed more than 20 response deadlines in either the preliminary phase or the main task, yielding 50 subjects remaining in our subsequent analyses.

#### Preliminary Phase

Before the main phase of the task subjects competed a preliminary task phase to gain familiarity with the task. Our version of the Simon task used blue and green circles as stimuli. The blue or green color was either associated with a left or right hand response (the ‘E’ or ‘I’ buttons on the keyboard). On each trial, a green or blue circle was presented on the left or right side of the screen (Figure 4A). The timing of events followed the timing of Experiment 1. In the version of the Simon task used here, 75% percent of trials were congruent—that is, the side on which the stimulus was presented matched the correct response hand. On the remaining 25% of trials, subjects needed to use stimulus color and fully ignore the stimulus side in order to respond correctly. Participants completed 120 preliminary trials.

**Figure 4.**
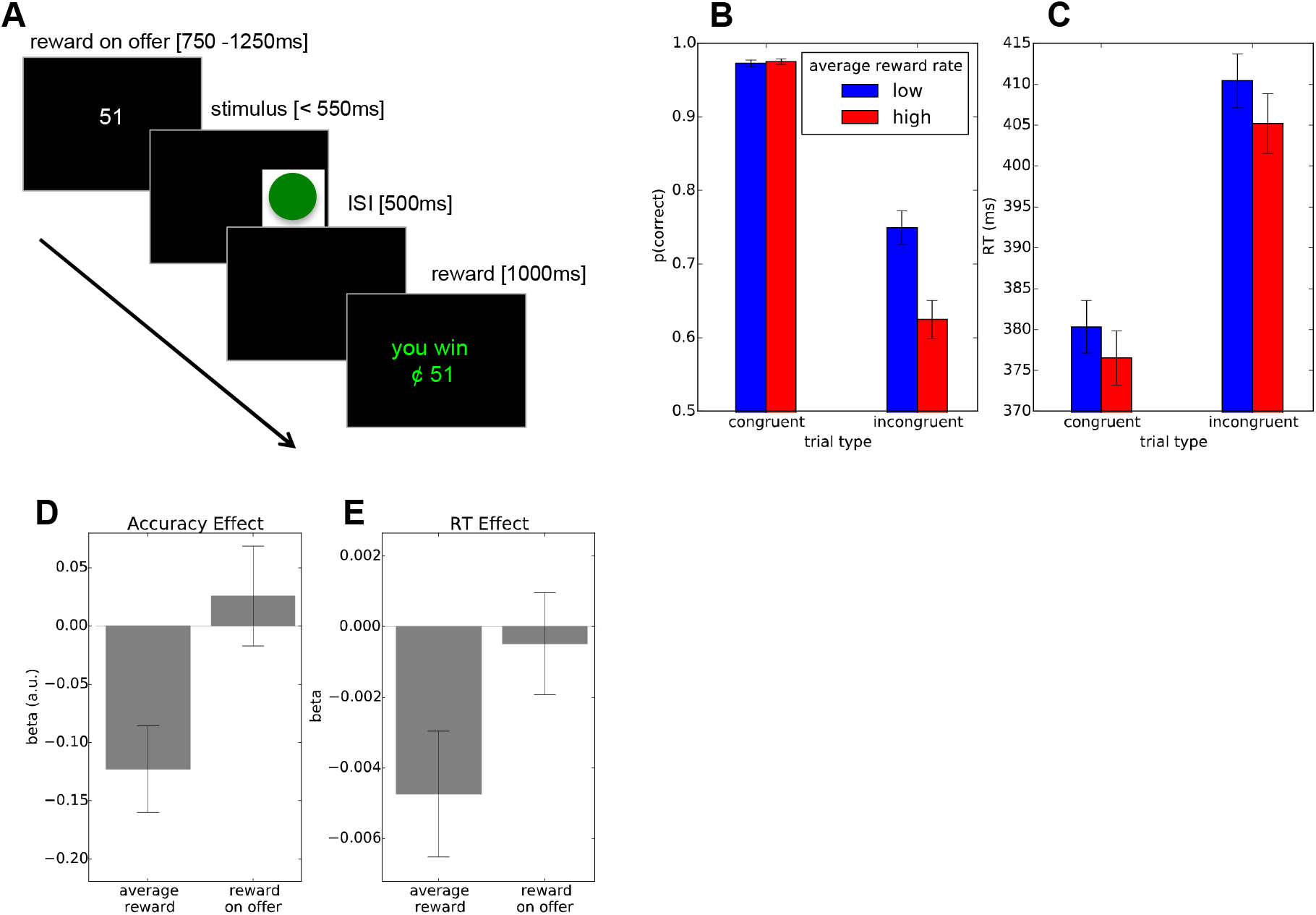
(A) Task flow in the Simon task. Before the stimulus is displayed, subjects are shown the potential reward for making a correct response, then they respond to a circle on the basis of its color, ignoring its location. (B-C) When the average reward rate was high, subjects made more errors on incongruent trials, where they needed to override innapropriate, prepotent responses, and made faster responses overall. Error bars indicate standard error of the mean. (D-E) Mixed-effects regressions revealed that the average reward rate significantly reduced accuracy and hastened RTs, but available reward did not exert an effect on either RT or accuracy. Error bars indicate standard error of the regression coefficient estimate.

#### Main Phase

Following the preliminary phase, subjects began the main phase of the experiment, following the same reward manipulation and timing as Experiment 1 (Figure 3A). Subjects completed as many trials as they could in 20 minutes.

#### Regression Analyses

To avoid bias issues stemming from simultaneously fitting *α* and estimating effects as a function of *α* (as described above), we simply used the best-fitting *α* of .0031 from Experiment 1 to calculate average reward rate. Mixed-effects regressions upon RTs were then conducted with the following terms:

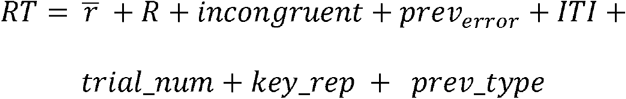

where *incongruent* codes whether the stimulus was incongruent or not, *stim_rep* represents whether the stimulus is a repetition from the previous trial, documented to facilitate faster RTs (Hommel, Proctor, & Vu, 2003), and *prev_error* and *prev_type* code for whether the subject committed an error on the previous trial and whether the previous trial’s stimulus was incongruent or not, which also exert effects on RTs and error rates (Ridderinkhof, 2002). Following Experiment 1, we also conducted a logistic regression using the same predictor variables with the exception of the ITI, with correct/incorrect response as the outcome variable.

### Results

In the Simon conflict task, subjects are required to make a right-button response to a green circle and a left-button response to a blue circle (Figure 4A). As the circle can appear either on the left or right side of the display, on most trials (“congruent” trials) subjects can effectively use the location of the stimulus to guide their responses, but on other trials (“incongruent” trials), subjects need to ignore the location of the stimulus in order to make a correct, color-based response. Because these congruent trials occurred 75% of the time, the more difficult incongruent trials require subjects to override a prepotent, stimulus-driven response established by congruent trials, and as a result, responses are markedly slower and more error-prone on these trials. As expected, we found that subjects made more errors (Figure 4B; (*β*= –3.029, SE=0.152, *p*<.0001) and were much slower to respond (Figure 4C; *β*=0.076, *SE*=0.003, *p*<.0001) on incongruent trials.

We then analyzed accuracy and RT as a function of average reward rate. Mirroring the results of the perceptual decision-making experiment, we found that a high average reward rate engendered an overall speeding of responses, as well as a marked decrease in accuracy on the more difficult incongruent trials. Mixed-effects regressions confirmed these effects statistically, finding main effects of average reward rate upon both accuracy and RT (Figures 4D and E; accuracy: *β*= 0.100, SE=0.036, *p*<.01; RT: *β*= −0.011, SE=0.001, *p*<.0001), but no effect of reward on offer (accuracy: *β*= 0.014, *SE*= 0.035, *p*= 0.686; RT: *β*= −0.001, SE=0.001, *p*=.615; see Tables 3 and 4). Taken together, these results again suggest that the opportunity cost of time may shift subjects’ speed-accuracy tradeoff on difficult trials, in favor of faster and less accurate responses.

**Table 3:** Mixed-effects Regression coefficients indicating the influence of the average reward rate and a number of other trial-by-trial covariates upon RTs in Experiment 2 (Simon Task). Asterisks denote significance at the .05 level.

| *Coefficient* | *Estimate (SE)* | *p-value* |
| --- | --- | --- |
| (Intercept) | 5.913 (0.009) | <.0001* |
| incongruent | 0.076 (0.003) | <.0001* |
| iti | -0.015 (0.002) | <.0001* |
| prev_errors | 0.010 (0.004) | 0.013* |
| trial_num | -0.005 (0.002) | 0.024* |
| key_rep | 0.017 (0.003) | <.0001* |
| prev_type | 0.020 (0.003) | <.0001* |
| reward | -0.001 (0.001) | 0.526 |
| **avg_reward** | **-0.006 (0.002)** | **0.002\*** |

**Table 4:** Mixed-effects logistic regression coefficients indicating the influence of the average reward rate and other trial-by-trial covariates upon accuracy in Experiment 2 (Simon Task). Asterisks denote significance at the .05 level.

| *Coefficient* | *Estimate (SE)* | *p-value* |
| --- | --- | --- |
| (Intercept) | 3.852 (0.154) | <.0001* |
| incongruent | -3.023 (0.152) | <.0001* |
| prev_errors | -0.214 (0.081) | 0.008* |
| trial_num | 0.107 (0.042) | 0.011* |
| key_rep | -0.068 (0.036) | 0.060 |
| prev_type | 0.151 (0.057) | 0.008* |
| reward | -0.037 (0.036) | 0.300 |
| **avg_reward** | **-0.106 (0.042)** | **0.012\*** |

Following previous work (Forstmann, van den Wildenberg, et al., 2008; Ridderinkhof, 2002), we visualized conditional accuracy functions—which plot accuracy as a function of RT quartile—revealing a marked speed-accuracy tradeoff on incongruent trials: faster responses were increasingly influenced by the irrelevant stimulus location (Figure 5A; RT accuracy effect: *β*= 1.132, SE=0.085, *p*<.0001). However, from the foregoing analyses it is not clear whether the effect of the opportunity cost of time is merely to move subjects *along* the SATF—from slower, accurate responses to faster, more error-prone responses—or if instead the average reward rate alters the SATF itself. As in the case of perceptual decision making, the first possibility suggests opportunity costs simply drive a speed-accuracy tradeoff over time investment; the second might reflect additional changes in the investment of cognitive resources per se.

**Figure 5.**
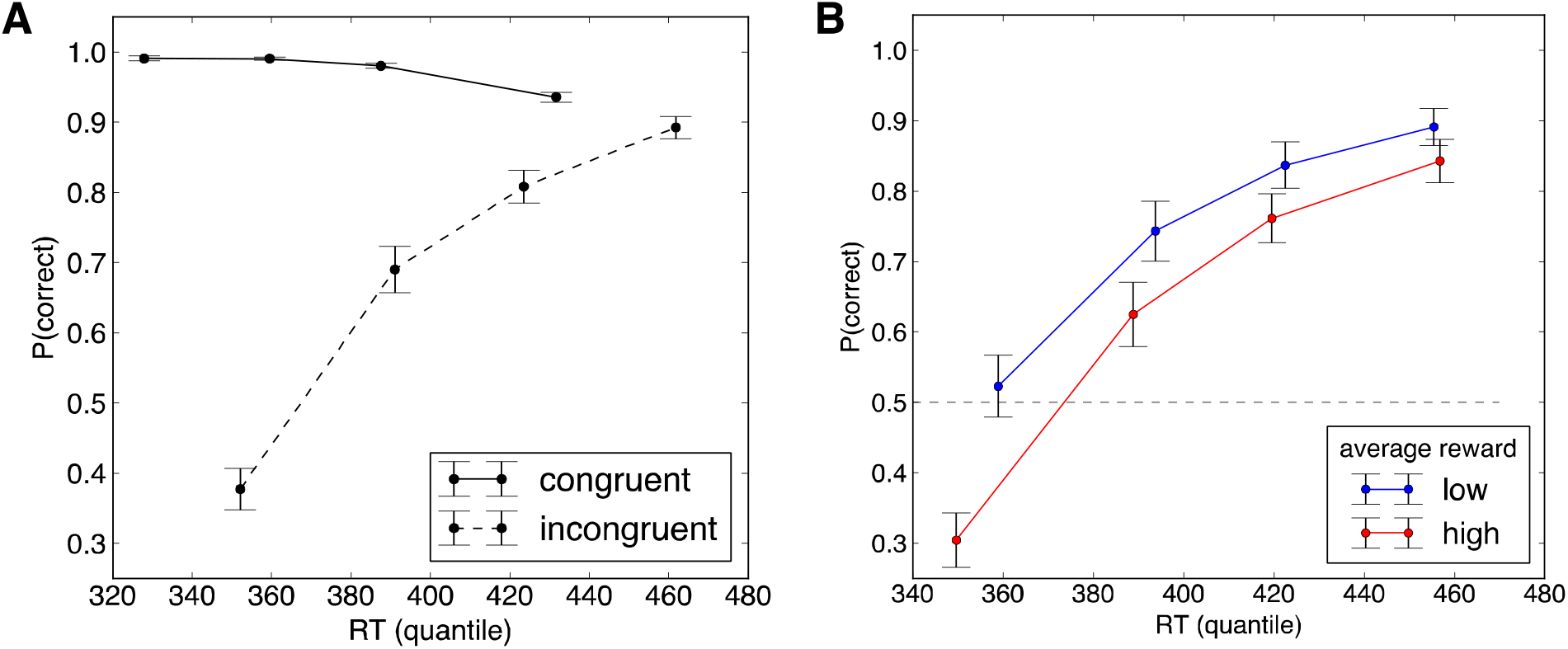
(A) Conditional accuracy plots reveal a marked speed-accuracy tradeoff on incongruent trials in the Simon task, such that fast responses appeared to be more driven by the task-irrelevant stimulus location, while slower responses yielded more accurate responses. (B) The opportunity cost of time altered the SATF on incongruent trials such that when the averag reward rate was high, responses were faster and appeared to be driven more by stimulus position. Error bars indicate standard error of the mean.

Visualized separately for high and low levels of the opportunity cost of time, the speed-accuracy tradeoff function (SATF) for incongruent trials (Figure 5B) indeed appears to shift downward, toward faster and more error-prone responses when the opportunity cost of time is higher. To quantify the change in SATF statistically, we used a mixed-effects logistic regression which jointly estimated the effect of average reward and response speed upon trial-by-trial accuracy, finding that the average reward rate significantly and negatively predicted accuracy over and above RT (3= −0.258, SE=0.058, *p*<.0001; Table 5). In other words, because these accuracy changes were not accounted for by response speed adjustments, these opportunity cost-induced changes in accuracy are not attributable to simple shift along a fixed SATF but rather change the SATF itself, again possibly reflecting changes in resource investment. Indeed, reward-related changes in speed-accuracy curves in response conflict tasks have been interpreted as reflecting effortful mobilization of attentional resources (Hübner & Schlösser, 2010).

**Table 5:**
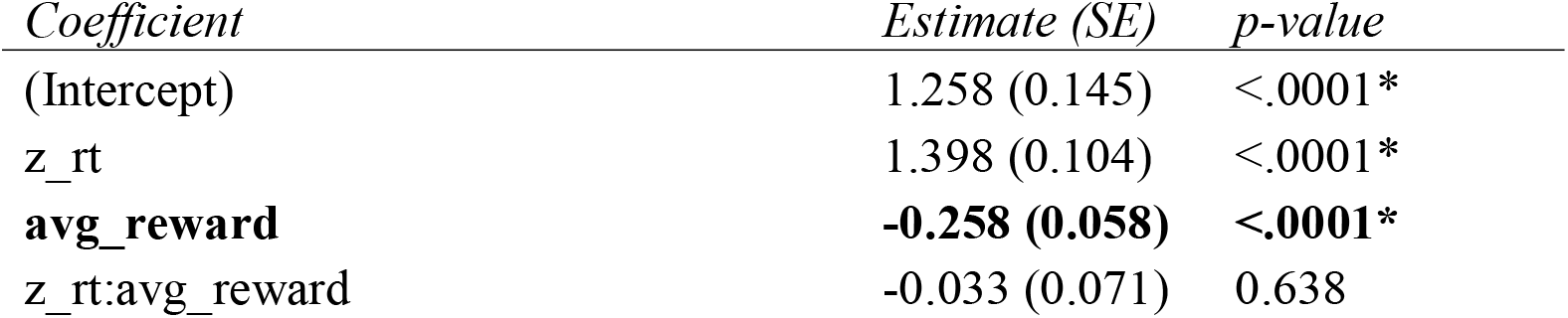
Mixed-effects logistic regression coefficients indicating the joint influence of the average reward rate and RT upon response accuracy in Experiment 2 (Simon Task, incongruent trials). Asterisks denote significance at the .05 level.

## Experiment 3: Task-Switching

We next sought to examine whether modulations of cognitive effort in accordance with the opportunity cost of time generalize beyond tasks or circumstances for which speed and accuracy objectively trade off—that is, situations where devoting additional time to making a response does not yield greater accuracy. By decoupling response speed and accuracy, changes in accuracy can be interpreted not as potentially strategic shifts in time allocation (as in the Simon Task) but potentially as more purely reflective of the amount of cognitive effort invested in a particular response. This would be expected, for instance, if the link between opportunity cost and effort investment were obligatory (e.g. reflexive or Pavlovian; Niv et al., 2007; Boureau et al., 2015) rather than learned online.

In contrast, two recent theories suggest that people learn to select cognitively expensive versus cognitively cheap ‘heuristic’ strategies on the basis of their learned costs and efficacy (Kool, Gershman, & Cushman, 2017; Lieder & Griffiths, 2017; Shenhav et al., 2017). In turn, the observed shifts in cognitive effort allocation in response to changing task circumstances are thought to reflect rational cost-benefit decisions at the aggregate level—most notably, the sacrifice of accuracy to improve the speed with which responses are made (Lieder & Griffiths, 2017). The absence of an inherent SATF in the task used below allows for a stringent test of whether individuals’ withholding of cognitive effort in the face of high opportunity costs are adaptively learned responses or if they simply result from a reflexive, Pavlovian response to high costs. Put another way, the observation of an opportunity cost-induced decrease in accuracy with no apparent upside in speed would suggest, compellingly, that the modulations of cognitive effort observed here are not the result of an adaptive learning mechanism per se.

In a third and final experiment we examine how the opportunity cost of time bears upon behavior in a task-switching paradigm, for which flexible responding in the face of shifts in stimulus-response rules demands central executive resources (Blain, Hollard, & Pessiglione, 2016; Kool et al., 2010; Monsell, 2003), and critically, there is no apparent SATF.

### Methods

#### Participants

We recruited 42 participants on AMT who paid a fixed amount ($5 USD) plus a bonus contingent on their decision task performance, ranging from $1-3 USD. We excluded the data of 9 subjects who missed more than 20 response deadlines in either the preliminary phase or the main task, yielding 33 subjects remaining in our subsequent analyses.

#### Preliminary Phase

Before the manipulation of opportunity costs phase, participants first completed a task-switching paradigm in the absence of rewards to gain familiarity with the task. On each trial, a box appeared on screen and participants’ needed either to report whether the box that appeared on screen was blue or orange (the “COLOR” task) or whether the box’s fill was solid or striped (the “PATTERN” task). Critically, the position of the box on the screen (lower half versus upper half, counterbalanced across subjects) indicated which subtask the subject was to perform. Across both subtasks, responses were either associated with a left- or right-hand button press (e.g. blue = left, orange = right; solid = left, striped=right), using the ‘E’ or ‘I’ buttons on the keyboard. Mappings of stimuli features to keys were counterbalanced across participants. Following Kool et al. (2010), the sequence of subtasks followed an m-sequence-based order, in which half the trials repeated the previous subtask. The trial timing was the same as in the calibration phase of Experiment 1 except that to accommodate the increased difficulty of this task, the response deadline was set to 800ms. Participants completed 120 preliminary trials.

#### Main Phase

Following the preliminary phase, subjects began the main phase of the experiment, following the same reward-on-offer manipulation and timing as Experiment 1, again with the exception of the response deadline of 800ms. Subjects completed as many trials as they could in 15 minutes.

#### Data Analyses

As in Experiment 2, we used the best-fitting *α* of .0031 from Experiment 1 to calculate average reward rate. For RT plots, RTs were z-scored within-subject to take into account. Mixed-effects regressions upon RTs were conducted with the following terms:

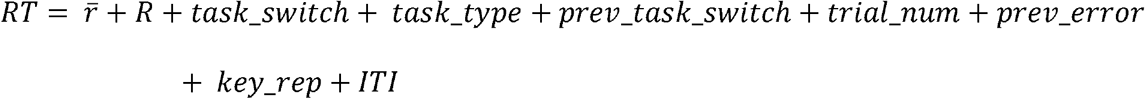

where *task_switch* codes whether the subtask repeated from the previous trial, *task_type indicated* whether the color versus pattern subtask was to be performed (thus capturing any differences in difficulty between subtasks). The accuracy analyses used a logistic regression using the same predictor variables with the exception of the ITI, with correct/incorrect response as the outcome variable.

### Results

In our task-switching paradigm, subjects responded to a stimulus based on a rule that varied from from trial to trial—here, subjects were required to indicate either the color (orange or blue) or the fill pattern (solid or striped) of a stimulus, depending on its position (Figure 6). On half of the trials, the required subtask (COLOR versus PATTERN) repeated, yielding a “repeat” trial, while the other half of trials entailed a switch to the other subtask (a “switch” trial). Following the previous experiments, we induced fluctuations in available reward (Figure 1C) and examined behavior as a function of the effective opportunity cost of time (i.e., the average reward rate).

**Figure 6.**
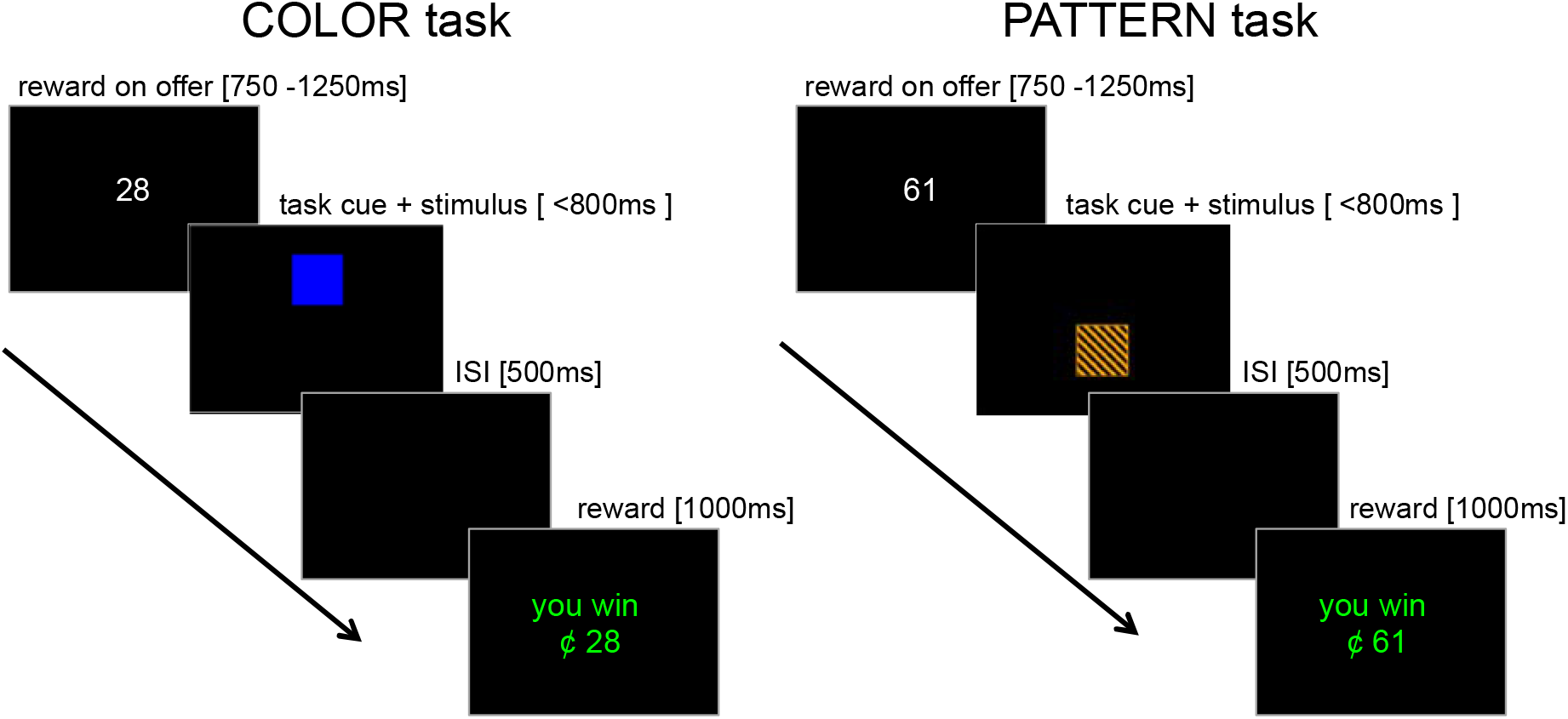
In the Task-Switching paradigm, subjects either indicated the color (blue or orange) or the pattern (stripes or solid) of a square, depending on its location on the display.

Mirroring previous work (Kool et al., 2010; Monsell, 2003), we found significant task switch costs in both response accuracy (Fig 7A; *β*= −0.258, SE=0.045, *p*<.0001) and RTs (Fig 7B; *β*=0.060, *SE* = 0.007)—task switches engendered less accurate and slower responses. Examining task-switching behavior as a function of the average reward rate, we found that the opportunity cost of time decreased accuracy, particularly on the more cognitively demanding switch trials and sped RTs overall. In other words, the apparent withdrawal of cognitive effort brought about by these opportunity costs manifested in an accuracy cost, particularly on the more cognitively demanding ‘switch’ trials. Mixed-effects regressions corroborated these effects (Figs 7C and D), revealing main effects of average reward rate upon both accuracy (*β*= −0.183, SE=0.051, *p*<.0001) and RT *β*=-0.009, SE=0.004, p<.05), and but again, no effects of reward-on-offer on either accuracy or RT (Tables 6 and 7).

**Figure 7.**
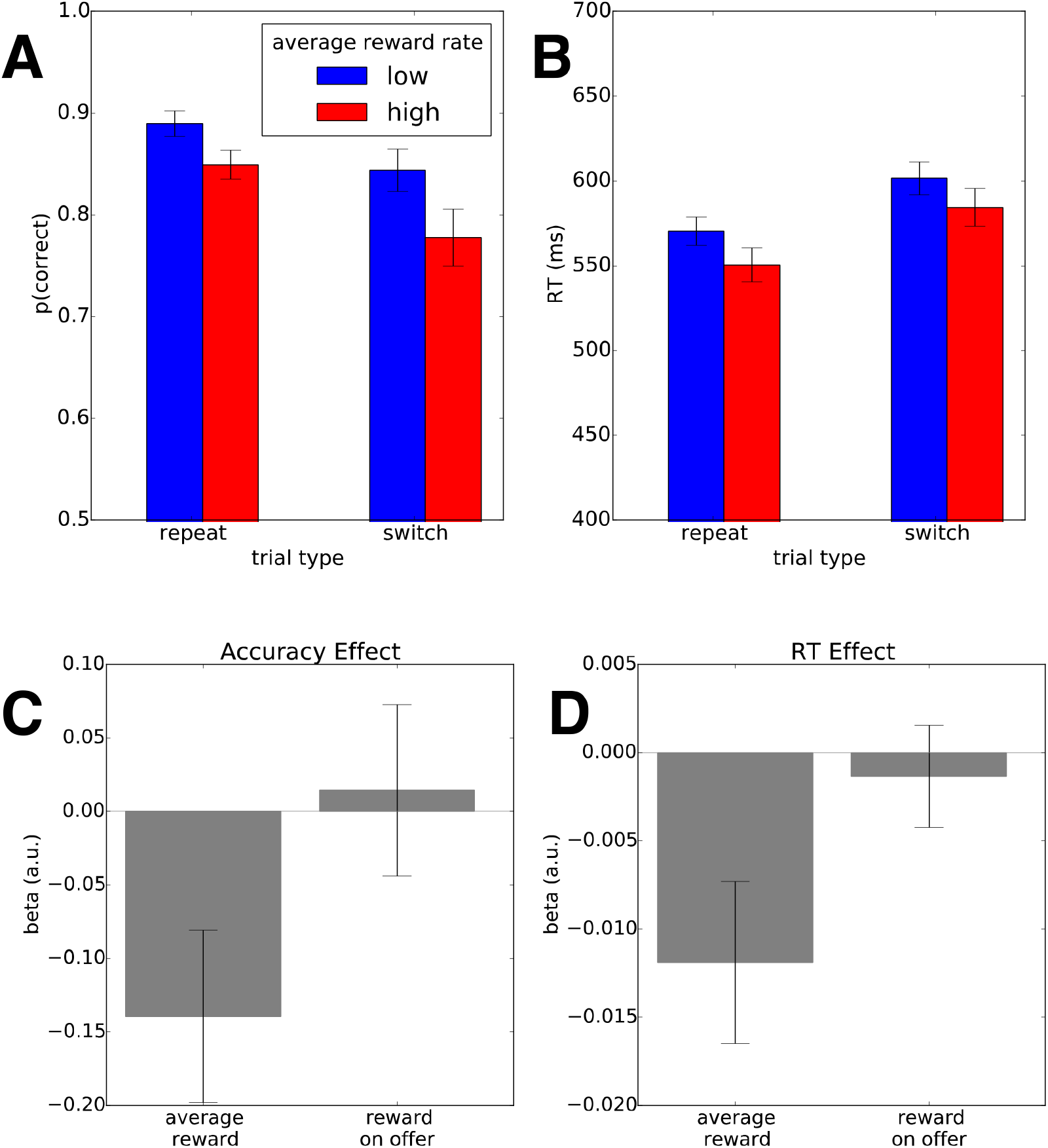
(A and B) Subjects exhibited significant task switch costs both in terms of accuracy and RT. When the average reward rate was high, subjects made more errors on task switches and made faster responses overall. Error bars indicate standard error of the mean. (C and D) Mixedeffects regressions revealed that on task switches, the average reward rate again significantly reduced accuracy and sped RTs, but available reward did not exert an effect on either RT or accuracy. Error bars indicate standard error of the regression coefficient estimate.

**Table 6:** Mixed-effects Regression coefficients indicating the influence of the average reward rate and a number of other trial-by-trial covariates upon RTs in Experiment 3 (Task-Switching). Asterisks denote significance at the .05 level.

| *Coefficient* | *Estimate (SE)* | *p-value* |
| --- | --- | --- |
| (Intercept) | 6.323 (0.016) | <.0001* |
| iti | -0.012 (0.002) | <.0001* |
| task_switch | 0.060 (0.008) | <.0001* |
| task_type | -0.009 (0.009) | 0.318 |
| prev_type | -0.011 (0.005) | 0.037* |
| trial_num | 0.006 (0.005) | 0.183 |
| prev_errors | 0.033 (0.008) | <.0001* |
| key_rep | -0.017 (0.005) | 0.002* |
| **avg_reward** | **-0.012 (0.004)** | **0.006\*** |
| reward | -0.001 (0.003) | 0.778 |

**Table 7:** Mixed-effects logistic regression coefficients indicating the influence of the average reward rate and a number of other trial-by-trial covariates upon accuracy in Experiment 3 (Task-Switching). Asterisks denote significance at the .05 level.

| *Coefficient* | *Estimate (SE)* | *p-value* |
| --- | --- | --- |
| (Intercept) | 1.779 (0.106) | <.0001* |
| task_switch | -0.268 (0.054) | <.0001* |
| task_type | 0.060 (0.083) | 0.469 |
| prev_type | -0.001 (0.047) | 0.979 |
| trial_num | 0.141 (0.058) | 0.014* |
| prev_errors | -0.065 (0.071) | 0.361 |
| key_rep | 0.069 (0.053) | 0.193 |
| **avg_reward** | **-0.120 (0.056)** | **0.031\*** |
| reward | 0.021 (0.046) | 0.650 |

Critically, we found no evidence for a positive SATF in overall behavior (RT effect on accuracy *β*= −0.17, SE=0.036, *p*<.0001, see Table 8)— faster RTs were associated with more accurate responding in both task switches and repetitions (Conditional Accuracy Functions plotted in Figures 8A and B). When we decompose the conditional accuracy function according to the observed average reward rates we find that, in both task switches and repetitions, we found that a high average reward rate simultaneously sped responding and decreased accuracy (Figures 8C and D), that is, holding the effects of reward rate constant, there is no evidence that the task trades off speed against accuracy. Interestingly, while the task-switching paradigm itself imposed no inherent speed-accuracy tradeoff—insofar as faster responses are also more likely to be correct, and therefore more rewarded, for a fixed reward rate— the opportunity cost of time exerted the same effects upon response speed and accuracy in the same way as observed in the previous experiment, further suggesting a decrease in effort investment.

**Figure 8:**
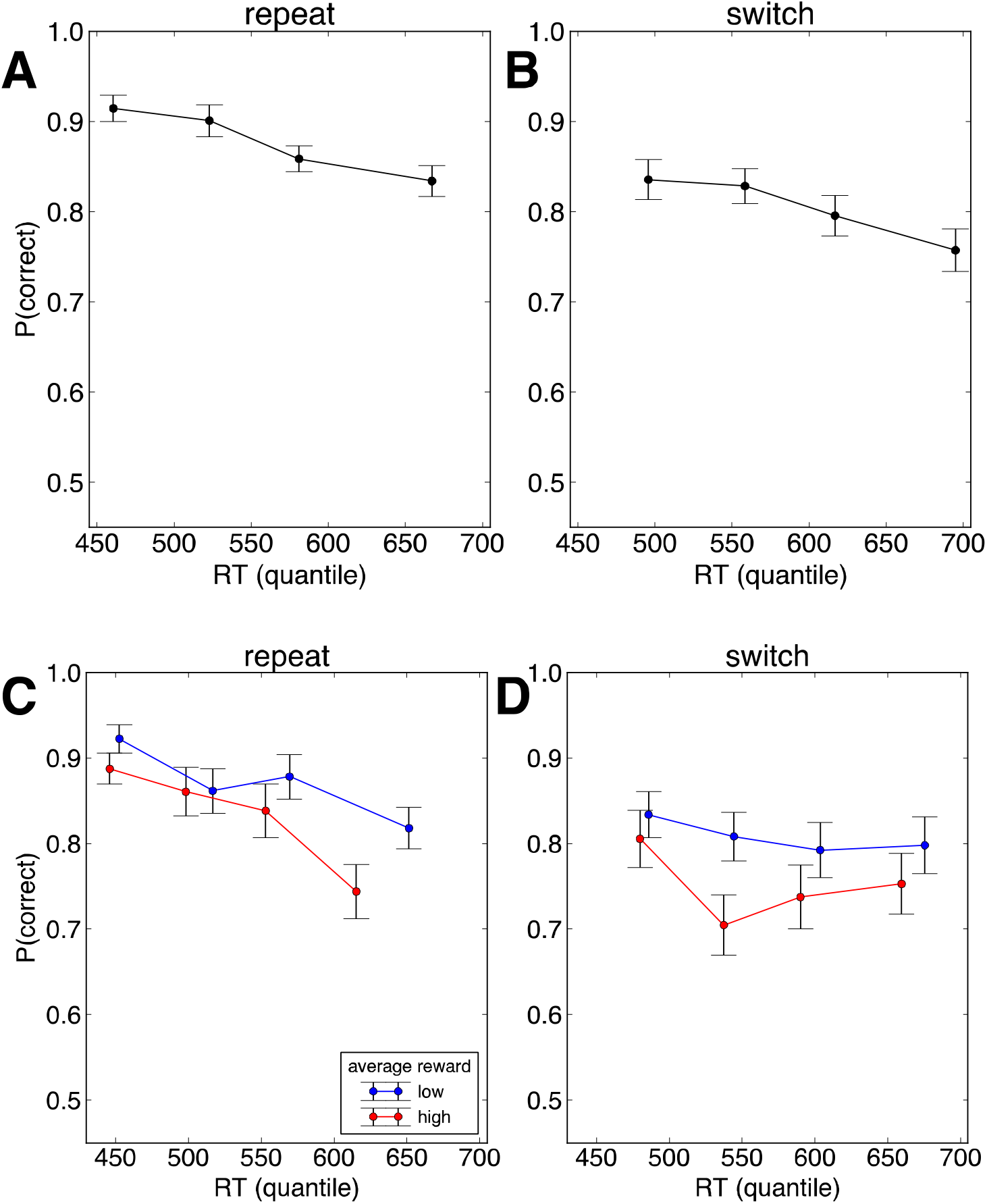
(A and B) Conditional accuracy plots reveal an absence of a positive speed-accuracy tradeoff on both task repetitions and task switch trials. (C and D) Conditional accuracy functions for stay and switch trials, split by average reward rate. Error bars indicate standard error of the mean.

**Table 8:** Mixed-effects logistic regression coefficients indicating the influence of RT in Experiment 3 (Task-switching). Asterisks denote significance at the .05 level. Note that this model specification does not include an intercept term because the combination of the task switch and task repeat terms is equivalent to the intercept term.

| *Coefficient* | *Estimate (SE)* | *p-value* |
| --- | --- | --- |
| task_switch | 1.292 (0.103) | <.0001* |
| task_repeat | 1.892 (0.096) | <.0001* |
| task_switch × z_rt | -0.123 (0.060) | 0.041* |
| task_repeat × z_rt | -0.295 (0.069) | <.0001* |

We also examined how individual differences in task-switching difficulty—thought to reflect processing efficiency and/or aversion to cognitive effort exertion (Kool et al., 2010)— predict the extent to which opportunity costs influence withholding of cognitive effort. Using separately obtained estimates of RT switch costs from a preliminary task phase, we found that greater RT switch costs predicted larger average reward rate-driven detriments to accuracy on switch trials (Figure 9). Critically, these separately obtained RT switch costs significantly predicted trial-by-trial accuracy modulations (*β* = 0.153, SE=0.065, *p*<05) even after controlling for 1) their predictive effect on individual’s overall error commissure rates and 2) individual differences in overall error commissure rates (Table 9). Put simply, individuals who experienced large task-switching costs were, in a subsequent task phase, more sensitive to the opportunity cost of time in their modulations of cognitive effort investment.

**Figure 9.**
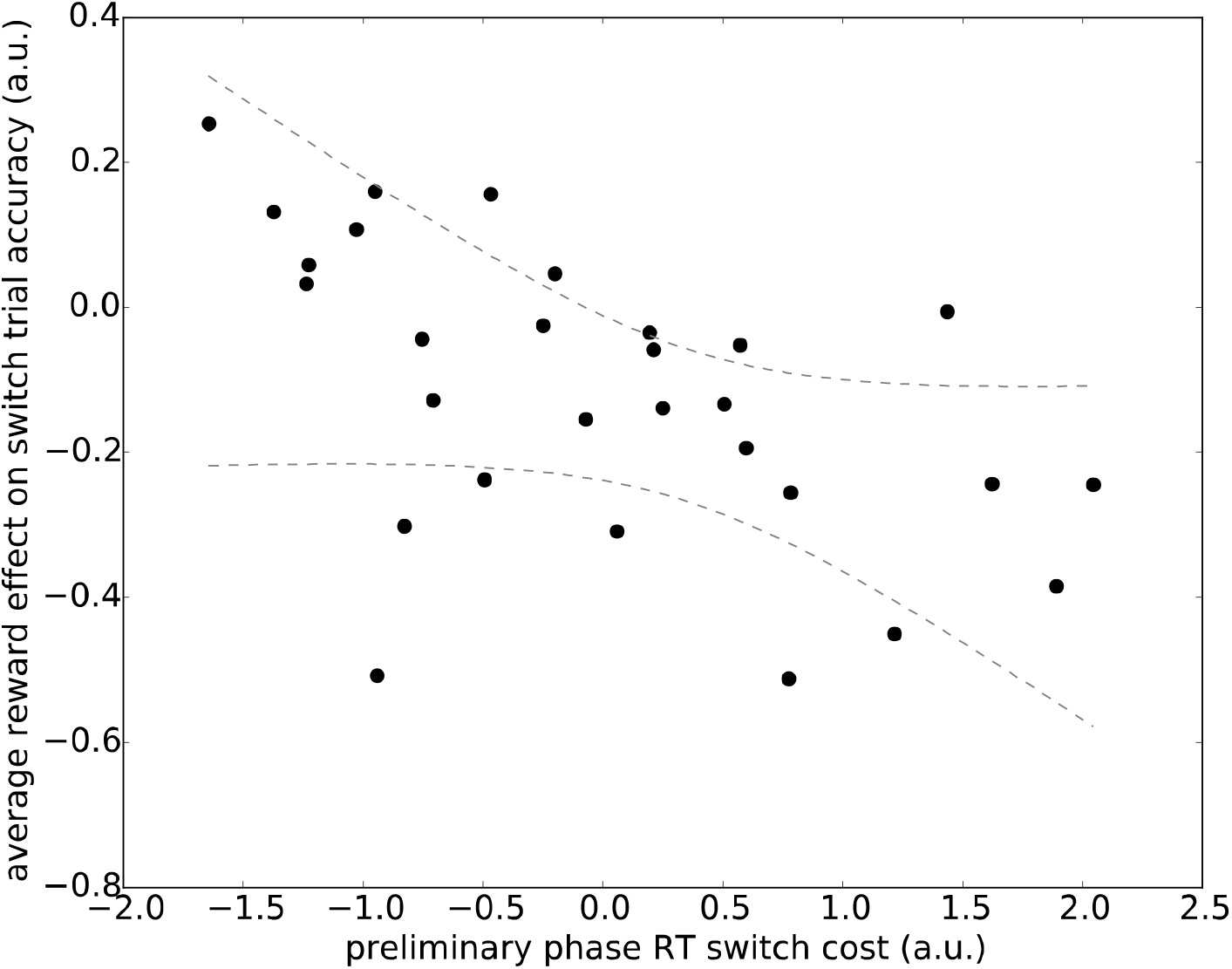
Task switch costs assessed in a preliminary phase (expressed in terms of RT) significantly predicted the effect of average reward upon task-switching accuracy in a subsequent phase. The regression lines is computed from the group-level fixed effect. Dashed lines indicate standard error about the regression line estimated from the group-level mixed effects regression.

**Table 9:** Mixed effects logistic regression coefficients indicating the joint influence of preliminary task switch costs (prelim_rt_switch_cost) and average reward effects (as well as other trial-by-trial covariates) upon accuracy in Experiment 3. Asterisks denote significance at the .05 level. The term of interest, depicted in bold, is the interaction between the preliminary task switch cost and the average reward rate (prelim_rt_switch_cost:avg_reward).

| *Coefficient* | *Estimate (SE)* | *p-value* |
| --- | --- | --- |
| (Intercept) | 1.704 (0.096) | <.0001* |
| z_prelim_rt_switch_cost | 0.114 (0.091) | 0.213 |
| trial_type | -0.268 (0.047) | <.0001* |
| prev_type | -0.013 (0.045) | 0.776 |
| prev_errors | -0.073 (0.070) | 0.302 |
| avg_reward | -0.127 (0.051) | 0.012* |
| reward | 0.013 (0.044) | 0.766 |
| z_prelim_rt_switch_cost × trial_type | -0.160 (0.045) | <.0001* |
| z_prelim_rt_switch_cost × prev_type | 0.072 (0.044) | 0.101 |
| z_prelim_rt_switch_cost × prev_errors | 0.034 (0.064) | 0.594 |
| **z_prelim_rt_switch_cost ×**<br>**avg_reward** | **-0.101 (0.049)** | **0.039\*** |
| z_prelim_rt_switch_cost × reward | -0.039 (0.043) | 0.362 |

## Discussion

Increasingly, researchers have attempted to extend accounts of rational decision making inward, to the question of how we allocate cognitive effort (Botvinick & Braver, 2015; Boureau et al., 2015; Kahneman, 2011; Kurzban et al., 2013; Shenhav et al., 2017). While the idea that cognitive effort should be invested (or withheld) in accordance with costs and benefits is intuitively appealing, it has been difficult to demonstrate conclusively. Here we leveraged research on the opportunity cost of time (Beierholm et al., 2013; Niv et al., 2007), which proposes a computationally explicit (though not necessarily exclusive) source for the costs of cognitive effort. Manipulating this quantity —operationalized as the average reward rate— allowed us to reveal how people withhold cognitively effortful processing when time is expensive, but will readily expend cognitive effort when time is ‘cheap’. By providing convergent evidence across diverse task domains, our approach yields a clear picture of how people negotiate the ubiquitous effort-reward trade-off.

We found that opportunity costs “invigorated” behavior such that individuals consistently sped their responses, mirroring past work (Beierholm et al., 2013; Guitart-Masip et al., 2011), and further, these opportunity costs resulted in more response errors. These accuracy effects could not simply be understood in terms of time allocation—rather, there were additional behavioral manifestations of the presumed withdrawal of cognitive effort in each of our three experiments. In a perceptual decision task, these costs reduced the amount of acceptable perceptual evidence required to make a decision (engendering faster, but less accurate decisions), but also altered the costly evidence accumulation process in favor of lower-fidelity perceptual evidence (bringing about less accurate decisions).

Similarly, in a response conflict task, the opportunity cost of time modulated the level of cognitive control that individuals applied to inhibiting inappropriate, prepotent responses—over and above what can be explained by a shift along a fixed speed-accuracy tradeoff (SATF). These results suggest that this moment-to-moment varying cost impacted strategic allocation of cognitive resources, perhaps in the same way that adjustments to cognitive control are made in response to individuals’ changing expectancy of response conflict (Gratton et al., 1992; Yu, Dayan, & Cohen, 2009). Intriguingly, this perspective suggests that the source of the performance costs seen in the face of response conflict (e.g., on incongruent trials) here and throughout the literature may in part be due to a withdrawal of effort on the basis of the added effort costs of responding correctly on those trials (rather than a direct effect of those trials being harder, per se). The observation the opportunity cost of time decreases performance when individuals experience response conflict lends credibility to the notion that failures of cognitive control, more generally, may arise in part from motivational deficits rather than cognitive ability.

In a final study, we observed that high opportunity costs prompted the withdrawal of cognitive effort during task-switching, which is understood to be cognitively demanding (Monsell, 2003). Notably, the speed-accuracy relationship implied by the task-switching paradigm (holding average reward fixed) does not appear to impose an inherent SATF upon behavior (in contrast to the Simon task; hence our choice of this task paradigm), and thus slower and presumably more effortful responses confer no advantages in terms of accuracy (and consequently, local rewards). However, individuals appeared to change their SATFs in response to the opportunity cost of time, simultaneously increasing speed and decreasing accuracy (Figure 8C and D), conceptually reproducing its observed effect in the Simon Task (Figure 5B) and suggesting a generality to the effects of the opportunity cost of time upon effort exertion— specifically, that opportunity costs appear to trigger a reflexive withdrawal of cognitive effort. Future investigation is needed to better understand how effort cost-benefit computations impact the relationship between speed and accuracy observed in many task domains.

Turning to individual differences, these modulations in effort expenditure depended in part on how costly task-switches are for an individual: when these cognitive costs loomed larger, individuals made larger opportunity-cost induced adjustments to effort allocation (Figure 9). This result dovetails well with the observation that individuals with fewer central executive resources avoid cognitive effort outlay compared to individuals with greater central executive function capacities (Kool et al., 2010; Otto, Skatova, Madlon-Kay, & Daw, 2015) and are more sensitive to shifts in benefits of effort expenditure (Sandra & Otto, 2018). To be sure, since the changes in accuracy observed here conferred no benefit in response speed (Figure 8), we interpret these effort adjustments as reflexive or obligatory. However, if such a reflexive strategy implements an approximation to adaptive cost-benefit tradeoff (albeit one not well suited to this task) it may still be adaptive in other circumstances for its strength to be modulated by an individual’s executive resources. The extent to which these opportunity cost-evoked effort modulations are governed by strategic or rational cost-benefit considerations (versus a reflexive response to costs) remains an important question for future research.

A potentially puzzling result uncovered across all three of our studies – which was also found in a similar opportunity cost manipulation (Beierholm et al., 2013; Guitart-Masip et al., 2011)—is the lack of significant effect of reward currently ‘on offer’ on effort investment. Indeed, even in estimating regressions that omit the average reward rate as a predictor, reward ‘on offer’ exerted no significant effect upon RT or accuracy in any of the three experiments reported (all ps > 0.25). Intriguingly, the motor control literature also reveals how available rewards can alter the speed-accuracy tradeoff—presumably by engendering increased cognitive resource investment—simultaneously engendering faster and more precise movements (Manohar et al., 2015). Taken together with the present work, a picture emerges that speed-accuracy tradeoffs are malleable and reflect, at any given moment, the investment (or withholding) of cognitive resources.

Opportunity cost models of time allocation models predict opposing effects of offered reward versus average reward: i.e., when the current trial’s reward is high, relative to the prevailing average, one should invest more time and resources. Indeed, much other work has found that various sorts of motivational cues—seemingly similar to the available reward ‘on offer’ manipulated here—increase engagement of executive or attentional resources (Bijleveld, Custers, & Aarts, 2010; Krebs, Boehler, & Woldorff, 2010; Manohar et al., 2015; Padmala & Pessoa, 2011). The reason for the lack of effect in the current paradigms remains unclear. It is possible that the average reward effect could contravene its presumably opposite effects on effort, or that the explicit numerical signaling of reward offers is less efficacious than that of obtained (also numerical) reward amounts.

Considering all three experiments together, the modulations of response accuracy revealed here suggest that the average reward rate signals not only a cost for slow physical movement—as originally ascribed to tonic dopamine level in the midbrain (Niv et al., 2007)— but also directs other, more cognitive speed-accuracy tradeoffs and the allocation of cognitive processing resources under the current circumstances. Indeed, the idea that the opportunity cost of time provides an internal signal that directs effortful processing away from the task at hand dovetails well with a previous account positing that mental effort is aversive because of opportunity costs inherent to the limited processing capacity of the central executive—i.e. the foregone benefits of putting these processing resources towards a different task (Kurzban et al., 2013). These results also hint that the apparent spontaneous fluctuations in cognitive effort outlay observed previously (Braver et al., 2003; Kahneman, 1973) could be explained, in part, by the opportunity cost of time.

Understanding how people modulate cognitive effort expenditures in accordance with opportunity costs is also relevant to neuroscientific research. Notably, there is compelling evidence that tonic midbrain dopamine level encodes the average reward rate of the environment (Beierholm et al., 2013; Niv et al., 2007) and this neuromodulatory system thus underpins an adaptive motivational control system. Relatedly, as neuroscientists find themselves increasingly interested in dopamine’s role in motivating deployment of cognitive resources during goal-directed behavior (Cools, 2015; Manohar et al., 2015; Westbrook & Braver, 2016), establishing the behavioral phenomenology of cognitive effort modulation will be critical for understanding the neuromodulator’s role in mobilizing cognitive resources.

## Acknowledgements

The authors are grateful to Claire Gillan, Peter Sokol-Hessner, Hanneke den Ouden, Andrew Westbrook, and Peter Dayan for helpful conversations during the development of this work as well as Marc Guitart-Masip for providing useful MATLAB code.

